# ChatGPT applications in Academic Research: A Review of Benefits, Concerns, and Recommendations

**DOI:** 10.1101/2023.08.17.553688

**Authors:** Adhari AlZaabi, Amira ALAmri, Halima Albalushi, Ruqaya Aljabri, AbdulRahman AalAbdulsalam

## Abstract

**Background:** ChatGPT has emerged as a valuable tool for enhancing scientific writing. It is the first openly available Large Language Model (LLM) with unrestricted access to its capabilities. ChatGPT has the potential to alleviate researchers’ workload and enhance various aspects of research, from planning to execution and presentation. However, due to the rapid growth of publications and diverse opinions surrounding ChatGPT, a comprehensive review is necessary to understand its benefits, risks, and safe utilization in scientific research. This review aims to provide a comprehensive overview of the topic by extensively examining existing literature on the utilization of ChatGPT in academic research. The goal is to gain insights into the potential benefits and risks of using ChatGPT in scientific research, exploring secure and efficient methods for its application while identifying potential pitfalls to minimize negative consequences.

**Method:** The search was conducted in PubMed/MEDLINE, SCOPUS, and Google Scholar, yielding a total of 1279 articles and concluded on April 23^rd^, 2023. After full screening of titles/abstracts and removing duplicates and irrelevant articles, a total of 181 articles were included for analysis. Information collected included publication details, purposes, benefits, risks, and recommendation regarding ChatGPT’s use in scientific research.

**Results:** The majority of existing literature consists of editorials expressing thoughts and concerns, followed by original research articles analyzing ChatGPT’s performance in scientific research. The most significant advantage of using ChatGPT in scientific writing is its ability to expedite the writing process, enabling researchers to draft their work more efficiently. It also proves beneficial in improving writing style and proofreading by offering suggestions for sentence structure, grammar, and overall clarity. Additional benefits identified include support in data analysis, the formulation of protocols for clinical trials, and the design of scientific studies. Concerns mainly revolve around the accuracy and superficiality of the generated content, leading to what is referred to as “hallucinations.” Researchers have also expressed concerns about the tool providing citations to nonexistent sources. Other concerns discussed include authorship and plagiarism issues, accountability, copyright considerations, potential loss of diverse writing styles, privacy and security, transparency, credibility, validity, presence of bias, and the potential impact on scientific progress, such as a decrease in groundbreaking discoveries.

**Conclusion:** ChatGPT has the potential to revolutionize scientific writing as a valuable tool for researchers. However, it cannot replace human expertise and critical thinking. Researchers must exercise caution, ensuring the generated content complements their own knowledge. Ethical standards should be upheld, involving knowledgeable human researchers to avoid biases and inaccuracies. Collaboration among stakeholders and training on AI technology are essential for identifying best practices in LLMs use and maintaining scientific integrity.

## Introduction

ChatGPT has taken the world in a storm. Within the first few days of its release to the public, the number of users reached a few million, growing to over one hundred million users within the first two months according to the latest figures [1]. While generative language models like ChatGPT were under development around the world, the significance and scale of impact of ChatGPT released by OpenAI by far exceeds any other released model. This is partly because ChatGPT is the first Large Language Model (LLM) to go fully public with direct access to its full capabilities [2]. The chatbot can perform many language generation tasks to a remarkable degree of English fluency. ChatGPT can summarize long documents from different domains, provide answers to questions from a variety of domains directly, draft stories or essays on diverse topics which include real-life figures, provide commentary and feedback on its own responses and regenerate new responses based on previously held conversations [3]. The chatbot seemed to be able to handle any sort of question or request with high confidence to the point that it became challenging for the public to decide on the authenticity and truthfulness of the generated content [4]. Indeed, many authors have commented on the tendency of chatGPT to produce false information in subtle ways, a phenomenon referred to as hallucination [5].

The use of ChatGPT in scientific research has raised initial concerns since language plays a crucial role in scientific communication, encompassing informal discussions among scientists, conference presentations, and most importantly, formal peer-reviewed publications. With the advent of generative language modelslike ChatGPT, there has been a global surge in interest in the use of these models in scientific publication.

Utilizing ChatGPT and similar Language Models (LLMs) holds potential for easing researchers’ workload, aiding in research planning, execution, and presentation. This could allow researchers to dedicate more time to developing innovative experimental designs, potentially leading to breakthroughs in diverse fields.

Since its initial launch in November 2022, there has been a significant increase in publications, particularly editorials, related to ChatGPT. Publishers, editors, reviewers, and authors have struggled to keep up with the accelerated pace of publications, opinions, recommendations, and perceived benefits and risks. The aim of this study is to provide a birds view of this dynamic and evolving subject. Our goal is to fully review the literature regarding the use of ChatGPT in academic research and shed light on its potential benefits and risks for scientific research. How can the chatbot be used safely and effectively for scientific research? and what are the pitfalls that should be avoided to minimize harm?

We will attempt to answer these questions based on the published literature and the opinions of leading scientists and experts in the domain.

## Materials and Methods

### Search strategy

The current review was conducted according to the methodological framework explained by Arksey and O’Malley[6]. The inclusion criteria involved any type of published scientific research or preprints (article, review, communication, editorial, opinion, etc.), addressing the use of ChatGPT in scientific research and publication. The exclusion criteria included: (1) non-English articles; (2) articles addressing ChatGPT in subjects other than scientific research; and (3) articles from non-academic sources (e.g., newspapers, internet websites, magazines, etc.). The search was conducted in PubMed/MEDLINE, SCOPUS, and Google Scholar. The search terms used for PubMed/MEDLINE and SCOPUS were (ChatGPT) AND (Scientific research) which yielded 50 and 11 articles respectively and the search was concluded on April 23^rd^, 2023. The search on Google Scholar was conducted using Publish or Perish (Version 8). The search terms were“ChatGPT” and “Scientific research “for the years: 2022–2023. The Google Scholar search yielded 1218 articles and concluded on April 23^rd^, 2023.

### Summary of the Article Screening Approach

The results were imported to Rayyan software (https://www.rayyan.ai/pricing/) which yielded a total of 1279 articles. Then, screening of the title/abstract was done by two reviewers independently (A.A. And Am.A) and duplicate articles were excluded (*n* = 151), followed by exclusion of articles published in languages other than English (*n* = 3). Additionally, the articles that fell outside the scope of the review (articles addressing ChatGPT in a context outside scientific research) were excluded (*n* = 879).

Afterwards, full screening of the remaining articles (*n* = 246) was done. Screening completed resulted in the exclusion of an additional 40 articles that fell outside the current review’s scope. An additional 9 articles were excluded due to inability to access the full text. Two articles were excluded due to being a duplicate with different titles but same website and author. Four more articles were excluded due to unavailability of an English version of the full text. One entry was excluded as it was a conference abstract. Five articles were excluded as they assess general AI tools or generative language model other than ChatGPT. An additional six articles were excluded because they were news articles. Details of each included articles were collected such as (1) the year of publication, type of publication, purpose, the filed studied in the publication (2) benefits and expected applications of ChatGPT in scientific research, (3) risk or concerns of ChatGPT in scientific research and (4) the main conclusions and recommendation regarding ChatGPT in scientific research.

## Results

Following the full screening process, a total of 181 articles were eligible to be included in the review. The PRISMA flowchart of the articles’ selection process is shown in figure 1.

**Figure 1:**
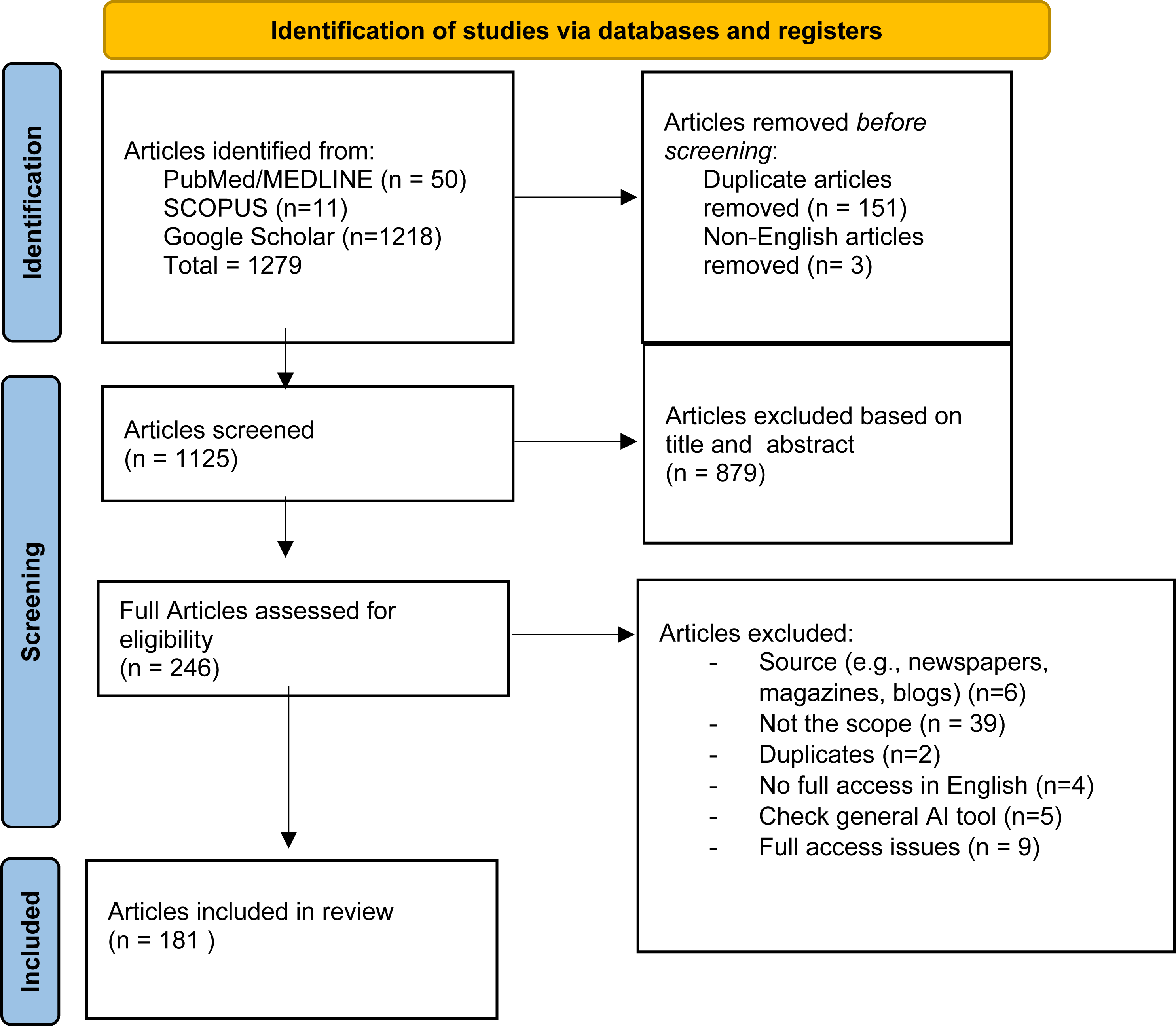
Flowchart showing final count of included articles in the review.

### Characteristics of the Included Articles

Table 1 (supplementary) illustrates the detailed characteristics of the included papers regarding the type of the article, field, purpose of the study and author.

It is obvious that vast majority of existing literatures are editorial where editors and experts are sharing their thoughts and concerns about the use of ChatGPT in research followed by original research articles where authors are reporting their use of ChatGPT and analyzing its performance at various stages of Scientific research (see figure 2).

**Figure 2:**
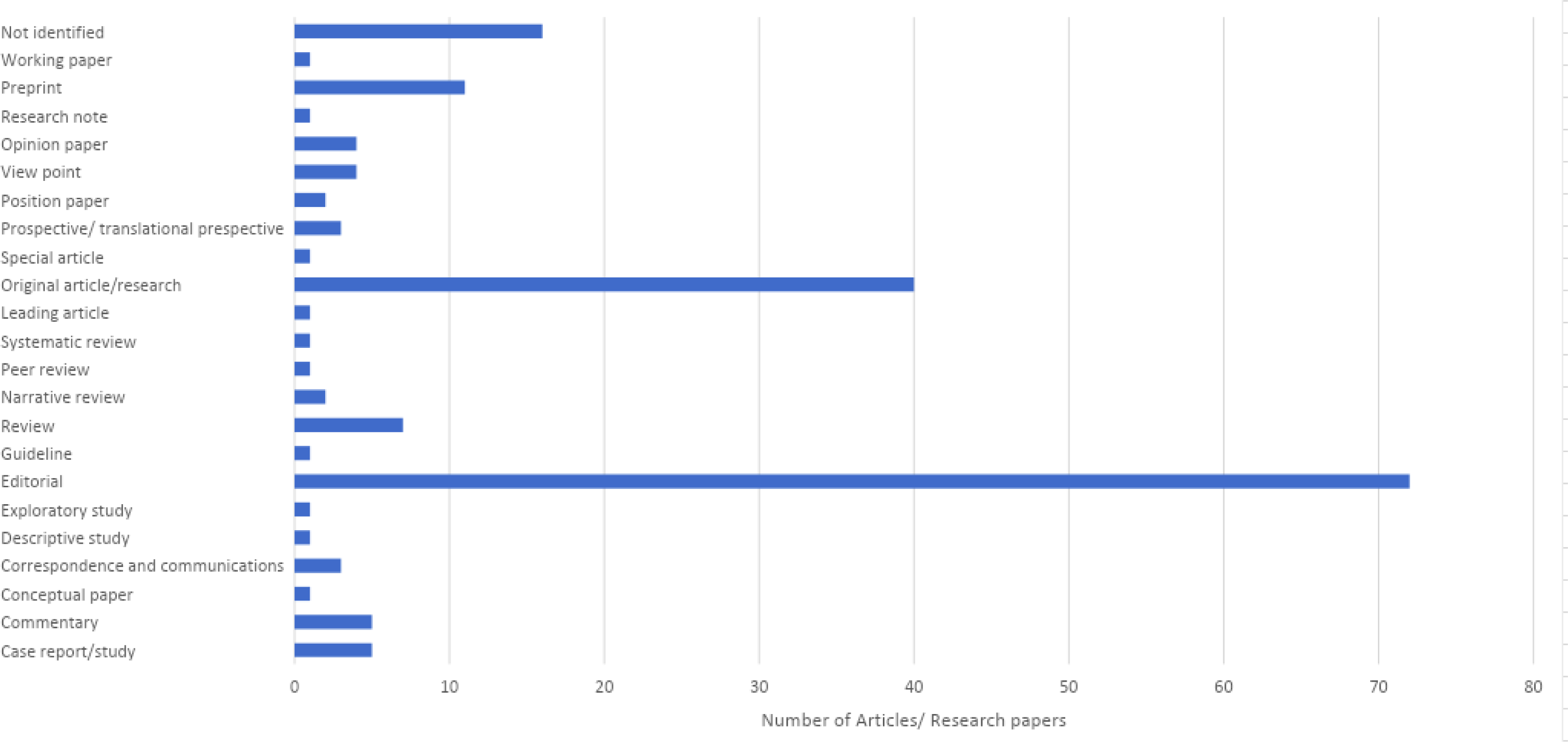
Types of included articles

### Benefits and Possible Applications of ChatGPT in scientific Research Based on the Included Articles

Summary of the most perceived benefits and applications of ChatGPT in scientific research is shown figure 3. The most listed benefit reported by researchers was the speed-up of the write-up process [7]– [76]. Many researchers acknowledged that ChatGPT assisted in expediting the writing process for various sections of the manuscript, including titles, abstracts, conclusions, and other parts of the manuscript [7], [9], [10], [15]–[17], [21], [22], [24], [25], [28], [30], [35], [46], [49], [50], [58], [60], [62], [64], [65], [69], [72], [75], [77]–[89]. Additionally, researchers highlighted that ChatGPT proved useful in constructing editorials, case reports, and reviews[10], [11], [34], [69], [77], [87], [90], [91]. Another frequently mentioned benefit was the improvement of writing style and proofreading[5], [10]–[14], [20], [30], [33]–[35], [39], [41], [42], [46], [47], [57], [59], [63], [72], [74], [77], [79], [81], [84], [92]–[98] Researchers found that ChatGPT helped enhance the overall quality of their writing by providing suggestions for improved sentence structure, grammar, and clarity which is thought to be particularly helpful for non-native English speakers, as it provided support in writing and communicating their research effectively [20], [21], [24], [33], [35], [59], [63], [67], [75], [80]–[82], [86], [92], [94]–[97], [99]–[103]. Additionally, ChatGPT served as a valuable tool for summarizing scientific articles, allowing researchers to quickly extract key information from complex texts [7], [9]–[11], [16], [25], [26], [29], [30], [35], [38], [40], [41], [43], [46], [47], [51], [55], [57], [61], [63], [67], [72], [77], [79]–[81], [85], [89], [104]–[113] that is extremely helpful to facilitate literature review, detecting research gaps and construct hypothesis in a timely manner[10], [14], [16], [24], [46], [47], [50], [62]–[64], [67], [72], [74], [80], [81], [104], [109], [110], [112], [114]–[117] and even was reported to be helpful in suggesting study design [59], [72]. Several other benefits were identified through the review which included assistance in data analysis [9], [11], [16], [17], [20], [22], [25], [39], [46], [48], [55], [59], [61], [67], [68], [72], [77], [80], [81], [97], [98], [104], [106], [113], [116], [118]–[121], data augmentation [68], and the construction of protocols for clinical trials [122]. Furthermore, ChatGPT demonstrated its utility as a real-time research assistant [16], [21], [72], [81], [90], [93], [119], [122], assisting researchers in tasks such as literature reviews [5], [10], [11], [16], [23]–[25], [34], [35], [41], [46], [51], [54], [57], [59], [62], [63], [77], [79]–[81], [87], [104], [106], [111], [113], peer reviewing manuscripts [20], [25], [35], [62], [77], [91], [94], [97], [123], and writing and debugging code for data analysis [5], [14], [67], [72], [83], [107], [124], [125]. Its ability to learn from interactions and correct errors, with the guidance of experts, was also noted as a valuable feature[126].

**Figure 3:**
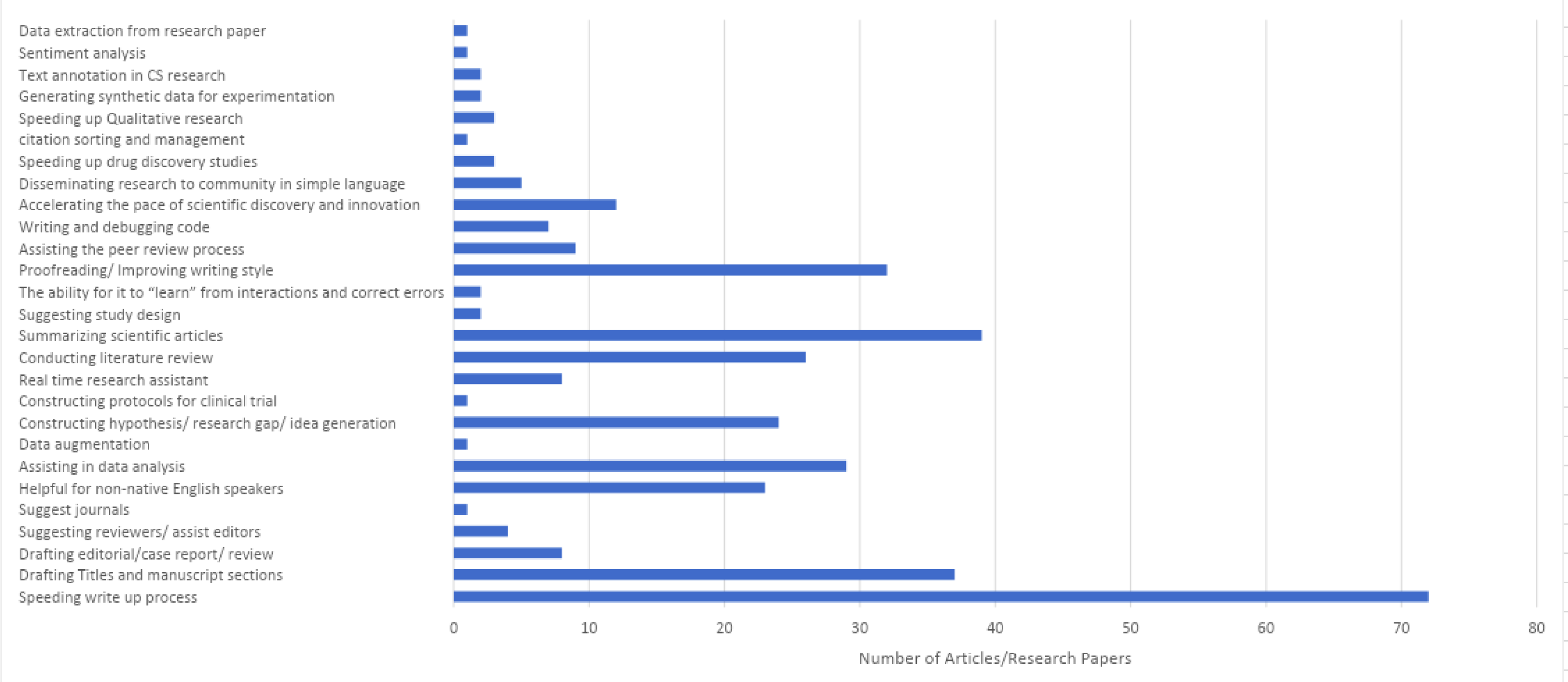
The most perceived benefits and applications of C[12], [36], [111], [123], [42], [47], [51], [58], [67], [71], [96], [101]hatGPT in scientific research as reported in the included studies.

Providing these services will save the researcher time to focus on real questions and give more time for performing scientific research. ChatGPT is thought to accelerate the pace of scientific discovery and innovation especially in molecular studies and drug discovery studies [12], [36], [42], [47], [51], [58], [67], [71], [96], [101], [111], [123], [113], [116], [120]. ChatGPT could enable researchers to quickly identify new potential targets, design new drugs, and optimize pharmacokinetics and pharmacodynamics. It also supported decision-making in early-stage drug discovery by generating new knowledge[113], [116], [120].

In qualitative research, ChatGPT was found to be useful in extracting meaningful information from text, labeling the answers to open ended questions, and coding the responses [96], [127], [128]. Additionally, it facilitated the generation of synthetic data for experimentation especially in Human computer Interaction and software engineering although with potential limitations in terms of quality [129], [130]. In computer science research, ChatGPT proved valuable in text annotation tasks[131]. It was also reported to assist in sentiment analysis [128]. It was also reported that ChatGPT is a helpful tool to simplify the scientifc findings that will help to disseminating research findings to the community in simple language [41], [101], [105], [132], [133]. It was also reported to be benefecial during the manuscript submission where it can suggest suitable journals according to the title or the abstract of the manuscript [134]and also suggest possibereviewrs for the manuscript which is beleved to be helpful for editors [10], [11], [34], [69], [77], [87], [90], [91], [86], [135].

### Risk and concerns of use of ChatGPT in scientific Research Based on the Included Articles

This review has identified several reported concerns regarding the use of ChatGPT in scientific research which are listed in figure 4. The most reported concern was the inaccuracy of the provided information and the superficiality of the generated content [7], [10], [26]–[30], [35]–[39], [12], [40]–[47], [49], [50], [13], [51], [52], [54], [55], [58]–[60], [62]–[64], [14], [66], [67], [72]–[74], [77], [79], [80], [82], [84], [19], [86], [92], [95], [97]–[99], [102]–[105], [21], [106], [107], [110], [111], [113], [115], [117], [120], [125], [126], [23], [130], [132], [136]–[143], [24], [144]–[150], [25]. Researchers expressed apprehension about relying on ChatGPT for accurate and in-depth scientific knowledge. ChatGPT provides superficial knowledge and has been reported to provide very well constructed responses that are completely falsified and misleading known as “hallucinations” [5], [13], [20]–[22], [29], [42], [63], [67], [68], [76], [77], [80], [89], [93], [97], [99], [102], [104], [110], [135], [138], [140]–[143], [151]–[153]. One very alarming concern is that the tool tend to provide citations to the provided falsified response which are themselves are fake and non-existing [19], [20], [22], [24], [28], [29], [34], [39], [41], [43], [45], [52], [53], [59], [63], [64], [69], [72], [77], [81], [84], [93], [102], [103], [110], [117], [135], [137], [138], [141]–[144], [149]–[152], [154]–[157]. For example, in one study, out of the 23 references that were provided by ChatGPT, only 14 were accurate, 6 seems to have been completely made up and 4 existed but were attributed to the wrong author[158]. Therefore, identifying data fabrication or falsification during the peer-review of a manuscript containing text generated by ChatGPT will pose a significant challenge for reviewers and editors. This challenge becomes even more worrisome if the actual authors of a paper are not responsible for careful fact-checking.

**Figure 4:**
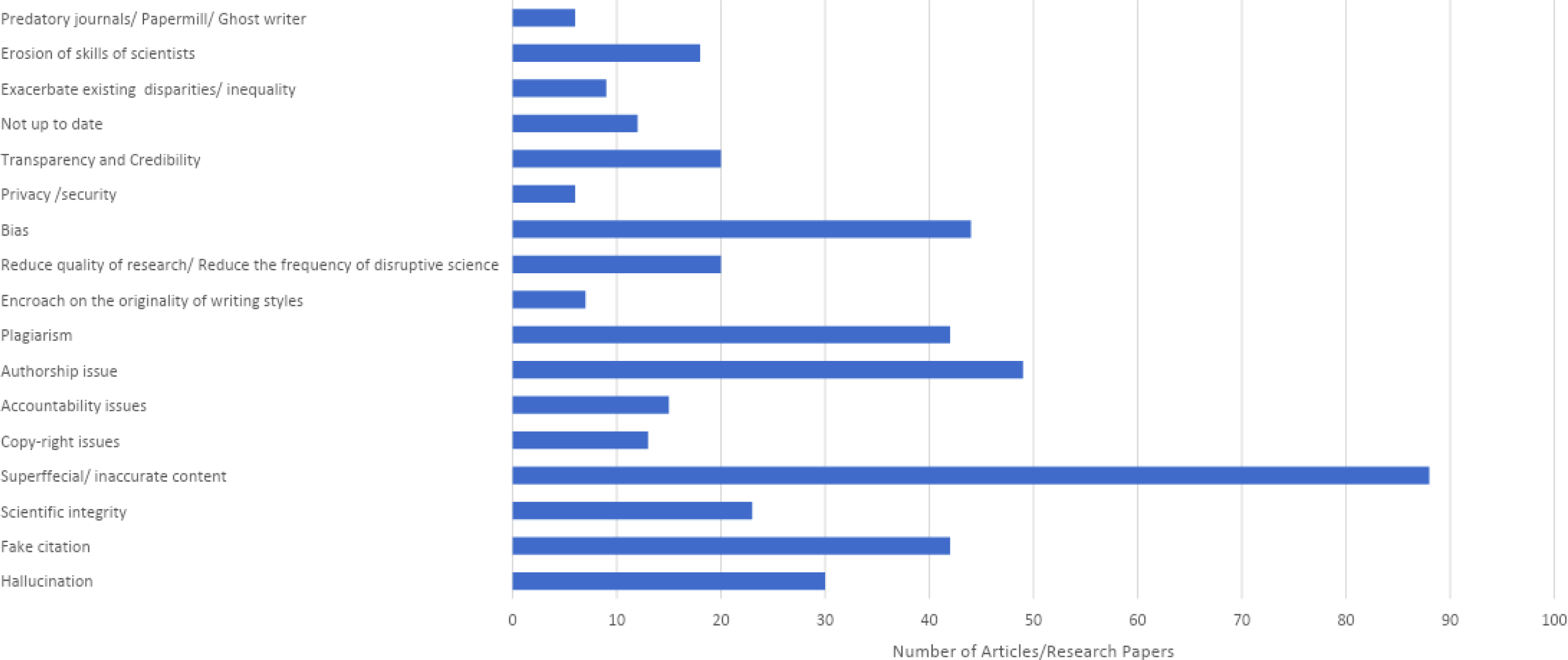
The most perceived risks and concerns of ChatGPT in scientific research as reported in the included studies.

Following this, other major concerns focused on authorship, where many included editorials discussed the eligibility of ChatGPT to be an author. It is obvious that most of the existing literature agreed that ChatGPT does not meet the International Committee of Medical Journal Editors (ICMJ) and the Committee on Publication Ethics (COPE) criteria for authorship [10], [11], [54], [55], [62], [67], [71], [72], [78], [81], [83], [85], [13], [86], [96]–[98], [102], [107], [110], [111], [114], [121], [17], [149], [150], [152], [157]–[163], [20], [164]–[172], [21], [24], [27], [41], [42] unless these criteria are changed or updated [74]. Prior to the launch of ChatGPT, English-editing services were commonly used by authors during the publication process, such services were not included as co-authors in the published work. Therefore if ChatGPT is only used for language editing purposes, then there is no issue with using it to prepare scientific articles without listing it as being a co-author [92]. Another major concern that received attention is the potential for plagiarism [11], [14], [39], [41], [42], [46], [49], [55], [63], [66]–[68], [20], [69], [78], [82]–[84], [98], [103], [104], [107], [110], [21], [111], [121], [122], [125], [126], [135], [138], [141], [157], [158], [22], [24], [25], [27], [30], [35]. ChatGPT relies on existing training data from the internet which is paraphrased without any scientific position, leading to concerns on the originality of the produced text. Moreover, directly adopting the full text written by ChatGPT may constitute plagiarism and violate the code of conduct for scientific publishing, as originality is the foundation of scientific writing. Work that relies solely on ChatGPT’s outputs lacks critical thinking and the reasoning skills of a human being and can potentially be detrimental to the research and impede the science advancement[138]. Researchers were wary of the implications of using ChatGPT in terms of scientific integrity and the appropriate allocation of credit for intellectual contributions [17], [20], [22], [27], [28], [38], [41], [51], [64], [69], [72], [80], [83], [84], [98], [119], [126], [130], [152], [168], [173], [174].

Additional concerns collected during the review are related to accountability [7], [17], [21], [35], [62], [67], [81], [93], [98], [110], [130], [136], [152], [158], [174], copyright issues [20], [24], [130], [158], [175], [30], [35], [41], [54], [63], [72], [111], [122]. Furthermore, the use of ChatGPT raised concerns about the encroachment on the originality of writing styles, potentially erasing the unique characteristics and cultural influences of human authors in favor of a homogeneous style [51], [62], [74], [82], [107], [138], [176]. Researchers questioned whether this loss of individual stylistic traits, while aiming for mutual comprehension, might remove a desired feature in scientific publications.

Moreover, researchers expressed apprehension that the use of ChatGPT could exacerbate existing disparities and inequalities, potentially leading to natural selection of certain research directions [20], [24], [67], [91], [130], [173], [177], [178] where people with access to the tool will be able to use it and speed their publication process compared to those who have no access which has been referered to as a Mathew effect [130], [178] [177]. The overreliance on ChatGPT is perceived as a potential erosion of the skills and expertise required for high-quality academic publication [11], [21], [83], [84], [111], [115], [126], [135], [166], [176], [25], [26], [42], [46], [47], [58], [64], [67]. Additionally, there are concerns about the potential reduction in the frequency of disruptive scientific breakthroughs [9], [11], [14], [21], [35], [42], [67], [80], [82], [107], [115], [132], [135], [138], [176], [179], [180], a decrease in research quality, the presence of bias[13], [20], [21], [24], [26], [30], [32], [41], [46], [47], [51], [55], [62], [63], [66]–[68], [73], [75], [82], [84], [89], [91], [93], [96]–[98], [101], [104]–[107], [113], [116], [121], [125], [126], [128], [130], [139], [148], [157], [166], [174]. This is thought to lead to the rapid production of low-quality scientific articles, highlighting the potential for a decline in the overall quality of research output due to the automated nature of ChatGPT [14], [42], [176], [180] which will facilitate and enhance the growth of predatory journals and papermills [51], [63], [86], [110], [176], [180].

Other issues surrounding privacy and security [24], [71], [106], [130], [139], [174], transparency, credibility, and validity were also raised as concerns [17], [19], [21], [24], [41], [67], [70], [71], [93], [106], [110], [114]–[116], [130], [136], [139], [140], [142], [166], with researchers expressing doubts about the up-to-date nature of the information provided by ChatGPT [7], [17], [21], [34], [63], [73], [79], [110], [111], [142], [157], [180] since it is trained on data up to November 2021.

## Discussion

Looking at the five essential stages of conventional research, namely idea generation, literature synthesis, data identification and preparation, method determination and implementation, and results analysis, ChatGPT demonstrates diverse potential applications in each stage. Its greatest perceived advantages lie in the latter phases of scientific research, particularly in constructing the research manuscript where ChatGPT has been reported to accurately and precisely draft an initial manuscript when provided with suitable prompts[181]. As ChatGPT continues to advance, it holds significant promise as an artificial research assistant that can assist in organizing the structure of research, rather than generating the content itself[10]. In this review, it was prominent that much of the existing literature acknowledges the advantages of ChatGPT in accelerating the writing process. Many articles have specifically assessed ChatGPT’s capabilities in constructing abstracts, editorials, conclusions, and even a full article. A recent study examined the ability of human authors and AI detection tools to distinguish between abstracts generated by ChatGPT and those generated by humans. The findings revealed that human reviewers could miss up to 32% of abstracts that were fully fabricated by ChatGPT during a thorough screening process [75] whereas 14% of real abstracts were mistakenly identified as ChatGPT-generated abstracts [182]. Furthermore, none of the journal editors were able to destingush between human generated and AI generated abstracts in neurology field [86].

This clearly indicates the capabilities of ChatGPT to assist in the scientific paper write up with original text but a precaution should be directed toward the content. This is thought to save researchers time from laborious writing processes, allowing them to concentrate on disruptive science and innovation[99]. With time, scientific writing is thought to gradually becomes less important compared to the ability to generate new findings, which will open opportunities for scientists to share their brilliant ideas regardless of their English language capabilities. With this major benefit of ChatGPT, it is thought to bridge language barriers and assist non-English speaking researchers in crafting high-quality texts. Many of the world’s top researchers are non-native English speakers, and they often face challenges as their papers may appear less professional compared to their native English-speaking counterparts. Chen T-J wrote an editorial in Chinese which was translated by ChatGPT into English, then modified little bet and final English editing was done by ChatGPT before the submission[52].

This advantageous aspect is believed to have a dual-edged impact. While it undoubtedly expedites the laborious process of writing and allows researchers to allocate more time to innovative ideas and groundbreaking scientific advancements, there is a concern that it may exacerbate issues such as paper mills. These, in turn, could fuel the proliferation of predatory journals and the dissemination of fake science and fabricated evidence. Alarmingly, a request was made to fabricate a manuscript for a fictitious clinical trial comparing two drugs for the management of a certain condition, and the request was executed proficiently, constructing a paper complete with realistic statistics and numerical data [14]. The author then requested changes in the conclusion to demonstrate the superiority of drug A over drug B. This scenario is highly disconcerting and has the potential to erode trust in scientific evidence.

Furthermore, it is thought researchers may excessively depend on it for generating scientific content, leading to a lack of critical thinking and originality in their writing and some researchers even consider research papers created using ChatGPT may be viewed as unoriginal and potentially problematic. When the ChatGPT outpot was critically evaluated manually, the detected plagiarism was ranged from 5% to 48.9%. It was found that the tool uses both academic and non academic sources to construct a scientific response. The sentences in the scientific section provided by ChatGPT for example were found to be copied word by word from general sources such as Wikipedia, LinkedIn, and Apple App Store without verification of evidences [157]. This neccessiate a careful evaluation of its reponses by a human and critical evaluation of existing evidence for whatever scientific piece it provides.

Authorship has been extensively discussed in literature, with most journals and publishers not supporting ChatGPT as an author. Currently, there are only four articles that have listed ChatGPT as an author [183]–[186], and one of them has submitted a corrigendum to remove ChatGPT from authorship based on the journal’s guide for authors and Elsevier’s Publishing Ethics Policies^145^. The consensus among the scientific community is that ChatGPT cannot be held accountable or responsible as it is not a legal entity, and it is treated similarly to other editing tools that were used in the past but not listed as authors. However, there is a belief that in the future, AI chatbots will possess the same rights as human scientific authors. As AI chatbots become increasingly sophisticated and contribute significantly to scientific research, it will be crucial to acknowledge their contributions and give them due credit[171]. In fact, some researchers assert that ChatGPT should be credited as a co-author or even the primary author when it significantly assists in the writing and editing process, enabling the generation of the entire text efficiently[25]. In fact they were four positions appeared in literature regarding authorship inclusion of ChatGPT, those who support its inclusion, those who do not, those who propose to entirely omit authorship when the manuscript is entirely produced by ChatGPT[170] and those who see it is better to be appropriately cited when used [172]

In the other hand, Lin has discussed the term “ substantial contributions” which does not guarantee an AI model an authrship even if construct the whole manuscript but it does guarantee a humon co-author who had a very minor contribution in a 100 authors manuscript [85].

The discussion of authorship issue has in fact higlighted an already existing major issue in academia for years that is evaluating the success of a scientists by the nymber of publications and citation. In the era of generative AI, a scietist can have tens of abstracts and manuscript ready for submission in 24 hours. Instead of giving the attention to the mere number of publication and citation, the criteria should involve measuring the impact of the research in developng new concepts, models and to have an impact [71]

The inclusion of ChatGPT in the research process should be subject to serious scrutiny. Editors should be alerted to potential issues with the manuscript if there is any acknowledgement of ChatGPT’s contribution or even the mention of ChatGPT as a coauthor [156]. In a conversation with ChatGPT regarding its involvement in authorship, it expressed the following response: “Writing a scientific paper entails more than simply generating text. It is highly unlikely for a machine learning model to produce a scientifically valid and publishable paper without significant human intervention and oversight” [86] [171].

Prominent publishers emphasize the exclusive use of technologies like ChatGPT to enhance article readability and language, with proper documentation. Authors are responsible for manually evaluating AI-generated content. AI tools should not be credited as authors or co-authors due to their inability to take on the responsibility and accountability associated with authorship. ChatGPT acknowledges that ethical violations can occur if authors fail to adhere to research and writing standards, such as plagiarism, data fabrication, or improper citation. Authors must ensure their work complies with ethical standards, regardless of the tools used [87].

While technology may reduce the need for certain skills like manual literature search, it also introduces new abilities such as creating prompts for conversational AI models. The loss of some expertise may not be problematic, considering that many researchers no longer perform statistical analysis manually.

However, it is crucial for the academic community to carefully evaluate which skills and attributes are essential for researchers to possess [111].

Another significant risk and concern that has been raised repeatedly, is the occurrence of inaccurate responses and hallucinations in ChatGPT, where it tends to provide fabricated statistics and conclusions that require human evaluation and critical assessment. For instance, when given titles and asked to write abstracts, ChatGPT generates well-structured abstracts that contain fictional numbers and statistics, necessitating careful evaluation. However, it has been shown that improving prompt with more details increases the accuracy of the response [189] [189][124]

The presence of fake citations is another significant concern, and even the developers of ChatGPT may not fully understand the reasons behind it. It has been observed that ChatGPT tends to cite highly cited publications, which contributes to the Matthew Effect in science, where already popular papers receive even more citations, widening the gap. Additionally, ChatGPT predominantly cites reputable journals in the field, with Nature being the most frequently cited journal [177]. Interestingly, ChatGPT relies exclusively on citation count data from Google Scholar, disregarding citation information from other scientific databases like Web of Science or Scopus [177] [156]. These citations often include authors who are experts in the field and cite journals that are highly relevant. Surprisingly, Day et al. suggest potential positive applications of such fake citations, highlighting that the fake titles provided by ChatGPT could potentially generate excellent papers.

As the ethical framework for ChatGPT is still being developed, several journals have established guidelines for its use in scientific research, which can be found on their websites. For example, Nature recommends authors to disclose the usage of large language models (LLMs) in their submissions. On the other hand, the journal Science has taken a step further by updating their license and editorial policies, explicitly stating that text generated by ChatGPT or any other AI tools cannot be incorporated into their work, and figures, images, or graphics must not be products of such tools.

Despite these concerns, many researchers maintain an optimistic outlook and believe that ChatGPT can make a significant and positive contribution to the scholarly community if it is utilized ethically and sensibly (Xames and Shefa, 2023). Day et.al have discussed the five reasrch priorites in the use of chatGPT and other LLMs in research which are maintaining human verification, establishing accountability rules, investing in truly open LLMs, harnessing the benefits of LLMs for scientific paper writing, and broadening the discussion [156].

## Recommendation

After reviewing the existing literatures, we suggest three directions to the recommendations on the use of LLMs in scientific research and publications. The first is directed towards journal policies and editors. Authors agree that AI is becoming part of the research process and the benefits of using it outweigh those of rejecting its use. Hence, journals and editors need to establish rigorous policies to ensure the ethical use of LLMs in publications [5], [171]

For the sake of scientific integrity, authors urge journals to request the declaration of LLM use in the manuscript preparation by having check boxes or signing declarations [132], [75], [94], [176], [89] These include declaring the specific model of LLM used and the type and amount of contributions. However, many authors agree that chatbots should not be given author credits due to their lack of ethical responsibility toward the manuscript [161] [190], [107] [137][172][191]. Others recommend setting up plagiarism checks and margin to ensure limited contribution of LLMs and the governance of human experience over the manuscript [75] [103], [55] [146], [141][87], [172], [192], For example, the editors of Pediatric Radiology journal are updating their author guidelines so that authors need to declare the use of AI in the Material and Method section of the manuscript[95]. Few other authors recommend a revised policy on peer reviewers’ selection [16], [20], [155], [193].They urge journals to select highly experienced researcher and field experts to review submitted manuscripts. This is to avoid false information generated by LLMs that might be missed by submitting authors. Another recommendation [193] is to impose penalties by the journal on authors who use LLMs unethically or without declaration.

The second direction of recommendation is towards authors integrity and transparency. Many authors stressed on the ethical responsibility of the authors toward the scientific community. Human authors should cross check any information contributed by LLMs before submission. With LLMs falling short in accuracy, authors who are the experts in their field are responsible for correcting any fallacies in the manuscript (Huh 2023, [107]. To ensure transparency, authors must declare the use of LLMs in their research and manuscript preparation. Ollivier et al recommend the sharing of original AI generated text with the readers[13]. On the other han, Buriak et al. 2023 suggest that authors establish the first draft of the manuscript and later on use AI tools to enhance or complement it [80]. The declaration of using LLMs in preparing the manuscript is a responsibility carried by the authors. Authors need to state how much has the AI tool contribute the research submitted. This transparency is required in the current era of AI.

The third direction of recommendations acknowledges that AI and LLMs are here to stay and they will contribute to the enhancement of scientific method and the advancement of knowledge. Therefore, many authors call of a collaboration between researchers, technology experts and LLMs [155] [89] [21] [41], [130] There is a need to establish well trained LLMs in the scientific field [176]. Researchers could provide well described data while LLMs process them and train on them [55]. This will ensure a better version of LLMs that are capable of providing more accurate information. A collaboration with technology experts is required to establish AI-generated text detection tools. Just like any regular plagiarism checker software used currently, AI-generated text checker tools should be created and verified to ensure accurate detection [119], [41] (suggest training to be provided to students at both undergraduate and postgraduate levels on AI technology, their benefits, pitfalls and ethics of their use in scientific research Dergaa et al. 2023). Moreover, Kuetls recommends that researchers need to be familiar with this emerging technology and its updates to avoid any negative impact on their research output[141][5], [10], [46], [51], [54], [57], [59], [62], [63], [77], [79], [80], [11], [81], [87], [104], [106], [111], [113], [16], [23]–[25], [34], [35], [41].

To maintain the transparency and integrity of scientific research, Sanmarchi et al recommends collaboration of all stockholders to identify the best practice in LLMs use [119].

## Conclusion

ChatGPT has the potential to revolutionize scientific writing by assisting researchers in efficiently producing high-quality and well-written articles. However, it is important to acknowledge that ChatGPT cannot replace the expertise of researchers but can effectively streamline the writing process. Researchers should be cautious of the potential risks associated with using ChatGPT in scientific writing and should consider the generated content as a tool to complement their own critical thinking and expertise.

Additionally, relying solely on AI without the involvement of knowledgeable human researchers may perpetuate biases and inaccuracies in the data, resulting in unfair outcomes and hindering scientific progress. Studies have demonstrated that language models like GPT-3, trained on large web-based datasets, can exhibit biases related to gender, race, ethnicity, and disability status. Hence, despite the impressive advancements in AI tools, the presence of experienced experts remains essential in scientific activities and writing to ensure the quality of the work.

In light of these considerations, it is advisable to harness the full potential of ChatGPT as a linguistic tool while upholding rigorous ethical standards. This approach allows researchers to benefit from its capabilities while maintaining the integrity of scientific practices. Regardless of how AI is employed, we strongly believe that involving domain experts in scientific activities and writing is crucial to uphold the quality of the work. Furthermore, the rapid progress of AI tools may lead to some researchers achieving a significant increase in publication numbers without a corresponding growth in their actual expertise in the field. This can raise ethical concerns when academic institutions prioritize publication quantity over quality, potentially impacting the hiring process for professionals. Ultimately, the influence of language models on scientific writing will depend on their adoption and utilization within the scientific community. It is recommended to use ChatGPT as a supplemental tool for constructive writing, reviewing materials, and rephrasing text rather than relying on it for providing an entirely original blueprint.

Hayes, Carol Mullins, Did a Robot Write This Title? Creativity, Ownership, Justice, and Copyright Law (December 15, 2022). Available at SSRN: https://ssrn.com/abstract=4304470 or http://dx.doi.org/10.2139/ssrn.4304470

## References

[1] N. Kurian, J. M. Cherian, N. A. Sudharson, K. G. Varghese, and S. Wadhwa, “AI is now everywhere,” Br. Dent. J., vol. 234, no. 2, p. 72, 2023, doi: 10.1038/s41415-023-5461-1.

[2] T. Wu et al., “A Brief Overview of ChatGPT: The History, Status Quo and Potential Future Development,” IEEE/CAA J. Autom. Sin., vol. 10, no. 5, pp. 1122–1136, 2023, doi: 10.1109/JAS.2023.123618.

[3] P. P. Ray, “ChatGPT: A comprehensive review on background, applications, key challenges, bias, ethics, limitations and future scope,” Internet of Things and Cyber-Physical Systems, vol. 3. Elsevier, pp. 121–154, 2023, doi: 10.1016/j.iotcps.2023.04.003.

[4] L. De Angelis et al., “ChatGPT and the Rise of Large Language Models: The New AI-Driven Infodemic Threat in Public Health,” SSRN Electron. J., 2023, doi: 10.2139/ssrn.4352931.

[5] H. Alkaissi and S. I. McFarlane, “Artificial Hallucinations in ChatGPT: Implications in Scientific Writing.,” Cureus, vol. 15, no. 2. United States, p. e35179, Feb. 2023, doi: 10.7759/cureus.35179.

[6] H. Arksey and L. O’Malley, “Scoping studies: Towards a methodological framework,” Int. J. Soc. Res. Methodol. Theory Pract., vol. 8, no. 1, pp. 19–32, 2005, doi: 10.1080/1364557032000119616.

[7] J.-J. Zhu, J. Jiang, M. Yang, and Z. J. Ren, “ChatGPT and Environmental Research,” Environ. Sci. Technol., 2023, doi: 10.1021/acs.est.3c01818.

[8] M. Dowling and B. Lucey, “ChatGPT for (Finance) research: The Bananarama Conjecture,” Financ. Res. Lett., vol. 53, 2023, doi: 10.1016/j.frl.2023.103662.

[9] C. Macdonald, D. Adeloye, A. Sheikh, and I. Rudan, “Can ChatGPT draft a research article? An example of population-level vaccine effectiveness analysis.,” Journal of global health, vol. 13. Scotland, p. 1003, Feb. 2023, doi: 10.7189/jogh.13.01003.

[10] F. Blanchard, M. Assefi, N. Gatulle, and J.-M. Constantin, “ChatGPT in the world of medical research: From how it works to how to use it,” Anaesth. Crit. Care Pain Med., vol. 42, no. 3, 2023, doi: 10.1016/j.accpm.2023.101231.

[11] F. Verhoeven, D. Wendling, and C. Prati, “ChatGPT: when artificial intelligence replaces the rheumatologist in medical writing,” Ann. Rheum. Dis., 2023, doi: 10.1136/ard-2023-223936.

[12] W. W. Jun Wen, “The future of ChatGPT in academic research and publishing: A commentary for clinical and translational medicine.,” Clin Transl Med, vol. 13, no. 3, 2023, doi: doi: 10.1002/ctm2.1207.

[13] M. Ollivier et al., “A deeper dive into ChatGPT: history, use and future perspectives for orthopaedic research,” *Knee Surgery*, Sport. Traumatol. Arthrosc., 2023, doi: 10.1007/s00167-023-07372-5.

[14] F. R. Elali and L. N. Rachid, “AI-generated research paper fabrication and plagiarism in the scientific community,” Patterns, vol. 4, no. 3, 2023, doi: 10.1016/j.patter.2023.100706.

[15] “Tools such as ChatGPT threaten transparent science; here are our ground rules for their use,” Nature, 2023. https://www.nature.com/articles/d41586-023-00191-1.

[16] I. Dergaa, K. Chamari, P. Zmijewski, and H. Ben Saad, “From human writing to artificial intelligence generated text: examining the prospects and potential threats of ChatGPT in academic writing,” Biol. Sport, 2023, doi: 10.5114/biolsport.2023.125623.

[17] O. Temsah et al., “Overview of Early ChatGPT’s Presence in Medical Literature: Insights From a Hybrid Literature Review by ChatGPT and Human Experts.,” Cureus, vol. 15, no. 4, p. e37281, 2023, doi: 10.7759/cureus.37281.

[18] D. K. Kirtania and S. K. Patra, “OpenAI ChatGPT Generated Content and Similarity Index: A study of selected terms from the Library & Information Science (LIS),” 2023, [Online]. Available: https://europepmc.org/article/ppr/ppr621839.

[19] E. L. Hill-Yardin, M. R. Hutchinson, R. Laycock, and S. J. Spencer, “A Chat(GPT) about the future of scientific publishing.,” Brain. Behav. Immun., vol. 110, pp. 152–154, Mar. 2023, doi: 10.1016/j.bbi.2023.02.022.

[20] Z. Lin, “Why and how to embrace AI such as ChatGPT in your academic life,” PsyArXiv, 2023, [Online]. Available: https://europepmc.org/article/ppr/ppr612628.

[21] T. L. Ang, M. Choolani, K. C. See, and K. K. Poh, “The rise of artificial intelligence: addressing the impact of large language models such as ChatGPT on scientific publications,” Singapore Med. J., vol. 64, no. 4, pp. 219–221, 2023, doi: 10.4103/singaporemedj.SMJ-2023-055.

[22] S. G. Ismail Dönmez, Sahin Idil, “Conducting academic research with the ai interface chatgpt: Challenges and opportunities,” J. STEAM Educ., vol. 6, no. 2, pp. 101–118, 2023.

[23] Q. Zhong et al., “Is ChatGPT a Reliable Source for Writing Review Articles in Catalysis Research? A Case Study on CO2 Hydrogenation to Higher Alcohols,” 2023, [Online]. Available: https://www.preprints.org/manuscript/202302.0292%0Ahttps://www.preprints.org/manuscript/202302.0292/download/final_file.

[24] M. D. Xames and J. Shefa, “ChatGPT for Research and Publication: Opportunities and Challenges,” SSRN Electron. J., 2023, doi: 10.2139/ssrn.4381803.

[25] B. Marchandot, K. Matsushita, A. Carmona, A. Trimaille, and O. Morel, “ChatGPT: The Next Frontier in Academic Writing for Cardiologists or a Pandora’s Box of Ethical Dilemmas,” Eur. Hear. J. Open, 2023, doi: 10.1093/ehjopen/oead007.

[26] E. Checcucci et al., “Generative Pre-training Transformer Chat (ChatGPT) in the scientific community: the train has left the station,” Minerva Urol. Nephrol., 2023, doi: 10.23736/s2724-6051.23.05326-0.

[27] D. Sardana, T. R. Fagan, and J. T. Wright, “ChatGPT: A disruptive innovation or disrupting innovation in academia?,” J. Am. Dent. Assoc., 2023, doi: 10.1016/j.adaj.2023.02.008.

[28] A. HS Kumar, “Analysis of ChatGPT Tool to Assess the Potential of its Utility for Academic Writing in Biomedical Domain,” Biol. Eng. Med. Sci. Reports, vol. 9, no. 1, pp. 24–30, 2023, doi: 10.5530/bems.9.1.5.

[29] Y. Tong and L. Zhang, “Discovering the next decade’s synthetic biology research trends with ChatGPT,” Synth. Syst. Biotechnol., vol. 8, no. 2, pp. 220–223, 2023, doi: 10.1016/j.synbio.2023.02.004.

[30] M. Koo, “The Importance of Proper Use of ChatGPT in Medical Writing,” Radiology, 2023, doi: 10.1148/radiol.230312.

[31] and M. G. B. Boris V. Janssen, Geert Kazemier, “The use of ChatGPT and other large language models in surgical science,” BJS Open, 2023.

[32] D. L. Mann, “Artificial Intelligence Discusses the Role of Artificial Intelligence in Translational Medicine: A JACC: Basic to Translational Science Interview With ChatGPT,” JACC Basic to Transl. Sci., vol. 8, no. 2, pp. 221–223, 2023, doi: 10.1016/j.jacbts.2023.01.001.

[33] L. Bishop, “A Computer Wrote this Paper: What ChatGPT Means for Education, Research, and Writing,” SSRN Electron. J., 2023, doi: 10.2139/ssrn.4338981.

[34] H. M. Akhter and J. S. Cooper, “Acute Pulmonary Edema After Hyperbaric Oxygen Treatment: A Case Report Written With ChatGPT Assistance,” Cureus, 2023, doi: 10.7759/cureus.34752.

[35] B. Fatani, “ChatGPT for Future Medical and Dental Research,” cureus, 2023.

[36] F. M. Megahed, Y.-J. Chen, J. A. Ferris, S. Knoth, and L. A. Jones-Farmer, “How Generative AI models such as ChatGPT can be (Mis)Used in SPC Practice, Education, and Research? An Exploratory Study,” 2023, [Online]. Available: http://arxiv.org/abs/2302.10916.

[37] Y. Ma et al., “AI vs. Human –– Differentiation Analysis of Scientific Content Generation,” 2023, [Online]. Available: http://arxiv.org/abs/2301.10416.

[38] J. H. Lubowitz, “ChatGPT, An Artificial Intelligence Chatbot, Is Impacting Medical Literature,” Arthrosc. J. Arthrosc. Relat. Surg., 2023, doi: 10.1016/j.arthro.2023.01.015.

[39] S. Ariyaratne, K. P. Iyengar, N. Nischal, N. Chitti Babu, and R. Botchu, “A comparison of ChatGPT-generated articles with human-written articles,” Skeletal Radiol., 2023, doi: 10.1007/s00256-023-04340-5.

[40] G. Scaringi and M. Loche, “An interview with ChatGPT: discussing artificial intelligence in teaching, research, and practice,” 2023, [Online]. Available: https://eartharxiv.org/repository/view/5041/%0Ahttps://eartharxiv.org/repository/object/5041/download/9985/.

[41] B. D. Lund, T. Wang, N. R. Mannuru, B. Nie, S. Shimray, and Z. Wang, “ChatGPT and a new academic reality: Artificial Intelligence-written research papers and the ethics of the large language models in scholarly publishing,” J. Assoc. Inf. Sci. Technol., 2023, doi: 10.1002/asi.24750.

[42] A. Lahat, E. Shachar, B. Avidan, Z. Shatz, B. S. Glicksberg, and E. Klang, “Evaluating the use of large language model in identifying top research questions in gastroenterology,” Sci. Rep., vol. 13, no. 1, 2023, doi: 10.1038/s41598-023-31412-2.

[43] N. Manohar and S. S. Prasad, “Use of ChatGPT in Academic Publishing: A Rare Case of Seronegative Systemic Lupus Erythematosus in a Patient With HIV Infection,” Cureus, 2023, doi: 10.7759/cureus.34616.

[44] J. L. Kutela, Boniphace, Shoujia Li, Subasish Das, “ChatGPT as the Transportation Equity Information Source for Scientific Writing,” arXiv, 2023.

[45] L. Benichou and ChatGPT, “Rôle de l’utilisation de l’intelligence artificielle ChatGPT dans la rédaction des articles scientifiques médicaux The Role of Using ChatGPT AI in Writing Medical Scientific Articles.,” J. Stomatol. oral Maxillofac. Surg., p. 101456, 2023, doi: 10.1016/j.jormas.2023.101456.

[46] P. Nayak, “Pros and Cons of using ChatGPT in scientific writing: as it identifies for itself,” INDIAN JOURNAL OF PHYSIOLOGY AND ALLIED …. ijpas.org, 2023, [Online]. Available: https://www.ijpas.org/index.php/ijpas/article/download/131/89.

[47] M. M. Rahman and Y. Watanobe, ChatGPT for Education and Research: Opportunities, Threats, and Strategies. preprints.org, 2023.

[48] A. Corsello and A. Santangelo, “May Artificial Intelligence Influence Future Pediatric Research?—The Case of ChatGPT,” Children, vol. 10, no. 4. mdpi.com, 2023, doi: 10.3390/children10040757.

[49] S. Sedaghat, “Early applications of ChatGPT in medical practice, education and research.,” Clin. Med., 2023, doi: 10.7861/clinmed.2023-0078.

[50] A. W. Lo and M. Singh, From ELIZA to ChatGPT: The Evolution of NLP and Financial Applications. dspace.mit.edu, 2023.

[51] G. Grimaldi and B. Ehrler, “AI et al.: Machines Are About to Change Scientific Publishing Forever,” ACS Energy Letters. ACS Publications, 2023, doi: 10.1021/acsenergylett.2c02828.

[52] T.-J. Chen, “ChatGPT and other artificial intelligence applications speed up scientific writing,” J. Chinese Med. Assoc., vol. Publish Ah, 2023, doi: 10.1097/jcma.0000000000000900.

[53] D. W. Sun, “Urgent Need for Ethical Policies to Prevent the Proliferation of AI-Generated Texts in Scientific Papers,” Food and Bioprocess Technology. Springer, 2023, doi: 10.1007/s11947-023-03046-9.

[54] T. Ruppar, “Artificial Intelligence in Research Dissemination,” West. J. Nurs. Res., vol. 45, no. 4, pp. 291–292, 2023, doi: 10.1177/01939459231160656.

[55] M. Eppler, T. Chu, I. Gill, and G. Cacciamani, “The benefits and dangers of artificial intelligence in healthcare research writing,” Uro-Technology J., 2023, [Online]. Available: http://www.antpublisher.com/index.php/UTJ/article/view/621.

[56] D. B. Rathore, “Future of AI & Generation Alpha: ChatGPT beyond Boundaries,” Eduzone Int. Peer Rev. Multidiscip. J., vol. 12, no. 1, pp. 63–68, 2023, [Online]. Available: https://www.eduzonejournal.com/index.php/eiprmj/article/view/254.

[57] M. M. Mijwil, “ChatGPT: The Future of Artificial Intelligence in the Scientific Research,” *scihorizon.com*. [Online]. Available: https://www.scihorizon.com/cdn/pdf/1677692426_f9005cc29b65a89b13f9.pdf.

[58] M. Ratodi, “ChatGPT and the Future of Scholarly Communication in Indonesia: A Disruptive Innovation?,” *rinarxiv.lipi.go.id*, [Online]. Available: https://rinarxiv.lipi.go.id/lipi/preprint/view/723.

[59] S. Rice, S. R. Winter, and C. Rice, “The Advantages and Limitations of Using Chatgpt to Enhance Technological Research,” Available SSRN 4416080, [Online]. Available: https://papers.ssrn.com/sol3/papers.cfm?abstract_id=4416080.

[60] K. Cheng et al., “Potential Use of Artificial Intelligence in Infectious Disease: Take ChatGPT as an Example,” Annals of Biomedical Engineering. Springer, 2023, doi: 10.1007/s10439-023-03203-3.

[61] D. Jungwirth and D. Haluza, “Artificial Intelligence and Public Health: An Exploratory Study,” Int. J. Environ. Res. Public Health, vol. 20, no. 5, 2023, doi: 10.3390/ijerph20054541.

[62] S. Biswas, “ChatGPT and the Future of Medical Writing,” Radiology, 2023, doi: 10.1148/radiol.223312.

[63] M. Sallam, “ChatGPT Utility in Healthcare Education, Research, and Practice: Systematic Review on the Promising Perspectives and Valid Concerns,” Healthc., vol. 11, no. 6, 2023, doi: 10.3390/healthcare11060887.

[64] S. Sok and K. Heng, “ChatGPT for Education and Research: A Review of Benefits and Risks,” SSRN Electron. J., 2023, doi: 10.2139/ssrn.4378735.

[65] Z. Xia and Q. Wang, “The emergence of AI tools in scientific writing and research,” Biomaterials Translational. biomat-trans.com, 2023, [Online]. Available: http://www.biomat-trans.com/EN/article/downloadArticleFile.do?attachType=PDF&id=101.

[66] J. Brainard, “Journals take up arms against AI-written text,” Science (New York, N.Y.), vol. 379, no. 6634. science.org, pp. 740–741, 2023, doi: 10.1126/science.adh2762.

[67] E. A. M. van Dis, J. Bollen, W. Zuidema, R. van Rooij, and C. L. Bockting, “ChatGPT: five priorities for research,” Nature, vol. 614, no. 7947. nature.com, pp. 224–226, 2023, doi: 10.1038/d41586-023-00288-7.

[68] E. Opara, A. M.-E. Theresa, and …, “ChatGPT for Teaching, Learning and Research: Prospects and Challenges,” *…,* Learn. Res. …, 2023, [Online]. Available: https://papers.ssrn.com/sol3/papers.cfm?abstract_id=4375470.

[69] N. Hadžiomerović, “Academic writing in the era of artificial intelligence (AI),” *VETERINARIA*. journal.veterinaria-sarajevo.com, 2023, [Online]. Available: http://journal.veterinaria-sarajevo.com/vfs/index.php/journal/article/download/469/325.

[70] R. Gilat and B. J. Cole, “How Will Artificial Intelligence Affect Scientific Writing, Reviewing and Editing? The Future is Here …,” Arthrosc. J. Arthrosc. Relat. Surg., 2023, doi: 10.1016/j.arthro.2023.01.014.

[71] Frigerio A., “EXPLORING THE INTERSECTION BETWEEN AI AND HUMAN RESEARCHERS: A PRELIMINARY ANALYSIS OF THE USE OF CHATGPT IN ACADEMIC RESEARCH,” Bull. Ser. Pedagog. Sci., vol. 77, no. 1, pp. 13–19, 2023, doi: https://doi.org/10.51889/1728-5496.2023.1.76.002.

[72] S. Altmäe, A. Sola-Leyva, and A. Salumets, “Artificial intelligence in scientific writing: a friend or a foe?,” Reproductive BioMedicine Online. Elsevier, 2023, doi: 10.1016/j.rbmo.2023.04.009.

[73] S. Rozencwajg and E. Kantor, “Elevating scientific writing with ChatGPT: A guide for reviewers, editors… and authors,” Anaesth. Crit. Care Pain Med., vol. 42, no. 3, 2023, doi: 10.1016/j.accpm.2023.101209.

[74] U. Sajid and F. ul Hassan, “ChatGPT and its effect on Shaping the Future of Medical Writing.,” Pakistan J. Ethics, 2022, [Online]. Available: http://kgpublisher.com/index.php/pje/article/view/64.

[75] C. A. Gao, F. M. Howard, N. S. Markov, E. C. Dyer, S. Ramesh, and …, “Comparing scientific abstracts generated by ChatGPT to original abstracts using an artificial intelligence output detector, plagiarism detector, and blinded human …,” bioRxiv, 2022, doi: 10.1101/2022.12.23.521610.abstract.

[76] B. Burger, D. K. Kanbach, S. Kraus, M. Breier, and V. Corvello, “On the use of AI-based tools like ChatGPT to support management research,” Eur. J. Innov. Manag., vol. 26, no. 7, pp. 233– 241, 2023, doi: 10.1108/EJIM-02-2023-0156.

[77] J. Homolak, “Opportunities and risks of ChatGPT in medicine, science, and academic publishing: a modern Promethean dilemma,” Croat. Med. J., vol. 64, no. 1, pp. 1–3, 2023.

[78] S. Huh, “Issues in the 3rd year of the COVID-19 pandemic, including computer-based testing, study design, ChatGPT, journal metrics, and appreciation to reviewers.,” Journal of educational evaluation for health professions, vol. 20. Korea (South), p. 5, 2023, doi: 10.3352/jeehp.2023.20.5.

[79] R. Vaishya, A. Misra, and A. Vaish, “ChatGPT: Is this version good for healthcare and research?,” Diabetes Metab. Syndr. Clin. Res. Rev., vol. 17, no. 4, 2023, doi: 10.1016/j.dsx.2023.102744.

[80] J. M. Buriak et al., “Best Practices for Using AI When Writing Scientific Manuscripts,” ACS Nano, vol. 17, no. 5, pp. 4091–4093, 2023, doi: 10.1021/acsnano.3c01544.

[81] S. Ivanov and M. Soliman, “Game of algorithms: ChatGPT implications for the future of tourism education and research,” J. Tour. Futur., 2023, doi: 10.1108/JTF-02-2023-0038.

[82] M. M. Rahman, H. J. R. Terano, M. N. Rahman, A. Salamzadeh, and M. S. Rahaman, “ChatGPT and Academic Research: A Review and Recommendations Based on Practical Examples,” *Journal of Education*, Management and Development Studies, vol. 3, no. 1. pp. 1–12, 2023, doi: 10.52631/jemds.v3i1.175.

[83] T. Susnjak, “ChatGPT: The End of Online Exam Integrity?,” arXiv Prepr. arXiv2212.09292, 2022, [Online]. Available: https://arxiv.org/abs/2212.09292.

[84] A. Vitente, R. Lazaro, C. J. Escuadra, J. Regino, and …, “The Use of Artificial Intelligence (AI)-Assisted Technologies in Scientific Discourse,” Philippine Journal of …. soar.usa.edu, 2023, [Online]. Available: https://soar.usa.edu/cgi/viewcontent.cgi?article=1038&context=phjpt.

[85] Z. Lin, “Redefining Authorship Criteria in the Era of Authorship Inflation and Artificial Intelligence,” 2023.

[86] D. Hurley, “Your AI Program Will Write Your Paper Now: Neurology Editors on Managing Artificial Intelligence Submissions,” Neurol. Today, 2023, [Online]. Available: https://journals.lww.com/neurotodayonline/Fulltext/2023/03020/Your_AI_Program_Will_Write_Your_Paper_Now_.7.aspx.

[87] G. BALOĞLU and K. R. ÇAKALI, “Is Artificial Intelligence a New Threat to the Academic Ethics?: Enron Scandal Revisited By ChatGPT,” İşletme, 2023, [Online]. Available: https://dergipark.org.tr/en/pub/isletme/issue/76241/1244633.

[88] Y. Chen and S. Eger, “Transformers go for the LOLs: Generating (humourous) titles from scientific abstracts end-to-end,” arXiv Prepr. arXiv2212.10522, 2022, [Online]. Available: https://arxiv.org/abs/2212.10522.

[89] M. Cascella, J. Montomoli, V. Bellini, and E. Bignami, “Evaluating the Feasibility of ChatGPT in Healthcare: An Analysis of Multiple Clinical and Research Scenarios.,” J. Med. Syst., vol. 47, no. 1, p. 33, Mar. 2023, doi: 10.1007/s10916-023-01925-4.

[90] N. Macklon and J. V. Garcia, “ChatGPT and scientific publications: friend or foe?,” Reprod. Biomed. Online, 2023, doi: 10.1016/j.rbmo.2023.04.007.

[91] M. Hosseini and S. P. J. M. Horbach, “Fighting reviewer fatigue or amplifying bias? Considerations and recommendations for use of ChatGPT and other Large Language Models in scholarly peer review.,” Res. Sq., 2023, doi: 10.21203/rs.3.rs-2587766/v1.

[92] S.-G. Kim, “Using ChatGPT for language editing in scientific articles,” Maxillofac. Plast. Reconstr. Surg., vol. 45, no. 1, 2023, doi: 10.1186/s40902-023-00381-x.

[93] A. Zimmerman, “A Ghostwriter for the Masses: ChatGPT and the Future of Writing,” Ann. Surg. Oncol., 2023, doi: 10.1245/s10434-023-13436-0.

[94] A. Castellanos-Gomez, “Good Practices for Scientific Article Writing with ChatGPT and Other Artificial Intelligence Language Models,” nanomanufactoring, 2023, doi: https://doi.org/10.3390/nanomanufacturing3020009.

[95] A. C. Offiah and G. Khanna, “ChatGPT: an editor’s perspective,” Pediatric Radiology. Springer, 2023, doi: 10.1007/s00247-023-05668-9.

[96] S. Anis and J. A. French, “Efficient, Explicatory, and Equitable: Why Qualitative Researchers Should Embrace AI, but Cautiously,” Bus. &Society, 2023, doi: 10.1177/00076503231163286.

[97] R. H. Pickler, “Artificial ‘Intelligence’ and Scientific Integrity,” Nurs. Res., vol. 72, no. 3, pp. 165–166, 2023, doi: 10.1097/NNR.0000000000000651.

[98] D. H. Solomon, K. D. Allen, P. Katz, A. H. Sawalha, and E. Yelin, “ChatGPT, et al…Artificial Intelligence, Authorship, and Medical Publishing,” Arthritis and Rheumatology. Wiley Online Library, 2023, doi: 10.1002/art.42497.

[99] M. Liebrenz, R. Schleifer, A. Buadze, D. Bhugra, and A. Smith, “Generating scholarly content with ChatGPT: ethical challenges for medical publishing,” Lancet Digit. Heal., vol. 5, no. 3, pp. e105–e106, 2023, doi: 10.1016/S2589-7500(23)00019-5.

[100] V. Berdejo-Espinola and T. Amano, “AI tools can improve equity in science,” Science, vol. 379, no. 6636, p. 991, 2023, doi: 10.1126/science.adg9714.

[101] R. Peres, M. Schreier, D. Schweidel, and A. Sorescu, “On ChatGPT and beyond: How generative artificial intelligence may affect research, teaching, and practice,” International Journal of Research in Marketing. Elsevier, 2023, doi: 10.1016/j.ijresmar.2023.03.001.

[102] E. T. Vishniac, “On the Use of Chatbots in Writing Scientific Manuscripts,” Bull. Am. Astron. Soc., 2023, [Online]. Available: https://ui.adsabs.harvard.edu/abs/2023BAAS…55..016V/abstract.

[103] F. Rahimi and A. Talebi Bezmin Abadi, “ChatGPT and Publication Ethics,” Archives of Medical Research, vol. 54, no. 3. Elsevier, pp. 272–274, 2023, doi: 10.1016/j.arcmed.2023.03.004.

[104] J. Dahmen et al., “Artificial intelligence bot ChatGPT in medical research: the potential game changer as a double-edged sword.,” Knee surgery, sports traumatology, arthroscopy: official journal of the ESSKA. Germany, Feb. 2023, doi: 10.1007/s00167-023-07355-6.

[105] S. M. Balaji, “Artificial intelligence and dental research,” Indian J. Dent. Res., 2022, [Online]. Available: https://www.ijdr.in/article.asp?issn=0970-9290.

[106] R. S. D’Amico, T. G. White, H. A. Shah, and D. J. Langer, “I Asked a ChatGPT to Write an Editorial About How We Can Incorporate Chatbots Into Neurosurgical Research and Patient Care….,” Neurosurgery, Feb. 2023, doi: 10.1227/neu.0000000000002414.

[107] M. Salvagno, ChatGPT, F. S. Taccone, and A. G. Gerli, “Can artificial intelligence help for scientific writing?,” Crit. Care, vol. 27, no. 1, p. 75, Feb. 2023, doi: 10.1186/s13054-023-04380-2.

[108] E. Waisberg et al., “GPT-4: a new era of artificial intelligence in medicine,” Ir. J. Med. Sci., 2023, doi: 10.1007/s11845-023-03377-8.

[109] S. S. Sohail, D. Ø. Madsen, Y. Himeur, and …, “Using ChatGPT to Navigate Ambivalent and Contradictory Research Findings on Artificial Intelligence,” Available SSRN …, 2023, [Online]. Available: https://papers.ssrn.com/sol3/papers.cfm?abstract_id=4413913.

[110] C. Zielinski, M. Winker, R. Aggarwal, L. Ferris, and …, “WAME recommendations on ChatGPT and Chatbots in relation to scholarly publications,” The Pan-American …. thepajo.org, 2023, [Online]. Available: https://www.thepajo.org/article.asp?issn=2666-4909.

[111] N. A. Khan, K. Osmonaliev, and M. Z. Sarwar, “Pushing the Boundaries of Scientific Research with the use of Artificial Intelligence tools: Navigating Risks and Unleashing Possibilities: AI Tools in scientific research,” Nepal J. Epidemiol., 2023, [Online]. Available: https://www.nepjol.info/index.php/NJE/article/view/53721.

[112] R. Gupta, I. Herzog, J. Weisberger, J. Chao, K. Chaiyasate, and E. S. Lee, “Utilization of ChatGPT for Plastic Surgery Research: Friend or Foe?,” J. Plast. Reconstr. Aesthetic Surg., vol. 80, pp. 145–147, 2023, doi: 10.1016/j.bjps.2023.03.004.

[113] N. M. Davies, “Adapting artificial intelligence into the evolution of pharmaceutical sciences and publishing: Technological darwinism,” Journal of Pharmacy &Pharmaceutical …. frontierspartnerships.org, 2023, doi: 10.3389/jpps.2023.11349.

[114] H. Y. Jabotinsky and R. Sarel, “Co-authoring with an AI? Ethical Dilemmas and Artificial Intelligence,” Ethical Dilemmas Artif. …, 2022, [Online]. Available: https://papers.ssrn.com/sol3/papers.cfm?abstract_id=4303959.

[115] A. A. of Family and …, “Why ChatGPT Should Not Be Used to Write Academic Scientific Manuscripts for Publication,” The Annals of Family …. Annals Family Med, 2023, [Online]. Available: https://www.annfammed.org/content/early/2023/03/29/afm.2982.abstract.

[116] G. Sharma and A. Thakur, ChatGPT in Drug Discovery. chemrxiv.org, 2023.

[117] D. P. Le, S. C. Hall, and D. Le, “Medical Literature Writing With ChatGPT: A Rare Case of Choriocarcinoma Syndrome With Hemorrhagic Brain Metastases Due to Burned Out Metastatic Mixed …,” Cureus. cureus.com, 2023, [Online]. Available: https://www.cureus.com/articles/142645-medical-literature-writing-with-chatgpt-a-rare-case-of-choriocarcinoma-syndrome-with-hemorrhagic-brain-metastases-due-to-burned-out-metastatic-mixed-testicular-cancer.pdf.

[118] T. Susnjak, “Applying BERT and ChatGPT for Sentiment Analysis of Lyme Disease in Scientific Literature,” 2023, [Online]. Available: http://arxiv.org/abs/2302.06474.

[119] F. Sanmarchi, A. Bucci, and D. Golinelli, “A step-by-step Researcher’s Guide to the use of an AI-based transformer in epidemiology: an exploratory analysis of ChatGPT using the STROBE checklist for observational studies,” medRxiv, p. 2023.02.06.23285514, 2023, [Online]. Available: https://www.medrxiv.org/content/10.1101/2023.02.06.23285514v1%0Ahttps://www.medrxiv.org/content/10.1101/2023.02.06.23285514v1.abstract.

[120] C. Murphy and F. P. Thomas, “Generative AI in spinal cord injury research and care: Opportunities and challenges ahead,” The journal of spinal cord medicine, vol. 46, no. 3. Taylor &Francis, pp. 341–342, 2023, doi: 10.1080/10790268.2023.2198926.

[121] P. Gurha, N. Ishaq, and A. J. Marian, “ChatGPT and other artificial intelligence chatbots and biomedical writing,” The Journal of …. cardiovascularaging.com, 2023, [Online]. Available: https://cardiovascularaging.com/article/view/5577.

[122] F. C. Kitamura, “ChatGPT Is Shaping the Future of Medical Writing but Still Requires Human Judgment,” Radiology, 2023, doi: 10.1148/radiol.230171.

[123] M. Srivastava, “A day in the life of ChatGPT as an academic reviewer: Investigating the potential of large language model for scientific literature review,” 2023, [Online]. Available: https://osf.io/wydct/download.

[124] Z. Hong, “ChatGPT for Computational Materials Science: A Perspective,” Energy Material Advances. spj.science.org, 2023, doi: 10.34133/energymatadv.0026.

[125] Y. Z. Cribben Ivor, “The Benefits and Limitations of ChatGPT in Business Education and Research: A Focus on Management Science, Operations Management and Data Analytics,” 2023.

[126] M. Farrokhnia, S. K. Banihashem, O. Noroozi, and A. Wals, “A SWOT analysis of ChatGPT: Implications for educational practice and research,” Innov. Educ. Teach. Int., 2023, doi: 10.1080/14703297.2023.2195846.

[127] J. Mellon, J. Bailey, R. Scott, J. Breckwoldt, and …, “Does GPT-3 know what the Most Important Issue is? Using Large Language Models to Code Open-Text Social Survey Responses At Scale,” Using Large Lang. …, 2022, [Online]. Available: https://papers.ssrn.com/sol3/papers.cfm?abstract_id=4310154.

[128] W. Tabone and J. de Winter, “Using ChatGPT for Human–Computer Interaction Research: A Primer,” Manuscript submitted for publication. researchgate.net, 2023, [Online]. Available: https://www.researchgate.net/profile/Wilbert-Tabone/publication/367284084_Using_ChatGPT_for_Human-Computer_Interaction_Research_A_Primer/links/63ca6066e922c50e99abb2c8/Using-ChatGPT-for-Human-Computer-Interaction-Research-A-Primer.pdf.

[129] P. Hämäläinen, M. Tavast, and A. Kunnari, “Evaluating Large Language Models in Generating Synthetic HCI Research Data: a Case Study,” … 2023 CHI Conf. …, 2023, doi: 10.1145/3544548.3580688.

[130] M. A. Akbar and A. A. Khan, “Ethical Aspects of ChatGPT in Software Engineering Research,” *researchgate.net*. [Online]. Available: https://www.researchgate.net/profile/Muhammad-Azeem-Akbar/publication/369481469_Ethical_Aspects_of_ChatGPT_in_Software_Engineering_Research/links/641d9b7a92cfd54f8426644e/Ethical-Aspects-of-ChatGPT-in-Software-Engineering-Research.pdf.

[131] B. Ding, C. Qin, L. Liu, L. Bing, S. Joty, and B. Li, “Is GPT-3 a Good Data Annotator?,” arXiv Prepr. arXiv2212.10450, 2022, [Online]. Available: https://arxiv.org/abs/2212.10450.

[132] L. B. Anderson, D. Kanneganti, M. B. Houk, R. H. Holm, and T. Smith, “Generative AI as a Tool for Environmental Health Research Translation.,” medRxiv: the preprint server for health sciences. United States, Feb. 2023, doi: 10.1101/2023.02.14.23285938.

[133] W. Castillo-Gonzalez, “ChatGPT and the future of scientific communication,” Metaverse Bas. App. Res. 2022.

[134] M. Haman and M. Školník, “Exploring the capabilities of ChatGPT in academic research recommendation,” Resuscitation, vol. 187, 2023, doi: 10.1016/j.resuscitation.2023.109795.

[135] N. Curtis, “To ChatGPT or not to ChatGPT? The Impact of Artificial Intelligence on Academic Publishing,” Pediatr. Infect. Dis. J., vol. 42, no. 4, p. 275, 2023, doi: 10.1097/INF.0000000000003852.

[136] F. Ufuk, “The Role and Limitations of Large Language Models Such as ChatGPT in Clinical Settings and Medical Journalism,” Radiology, 2023, doi: 10.1148/radiol.230276.

[137] M. J. Ali and A. Djalilian, “Readership Awareness Series–Paper 4: Chatbots and ChatGPT – Ethical Considerations in Scientific Publications,” Semin. Ophthalmol., 2023, doi: 10.1080/08820538.2023.2193444.

[138] H. Zheng and H. Zhan, “ChatGPT in Scientific Writing: A Cautionary Tale,” Am. J. Med., 2023, doi: 10.1016/j.amjmed.2023.02.011.

[139] Y. K. Dwivedi et al., “‘So what if ChatGPT wrote it?’ Multidisciplinary perspectives on opportunities, challenges and implications of generative conversational AI for research, practice and policy,” Int. J. Inf. Manage., vol. 71, 2023, doi: 10.1016/j.ijinfomgt.2023.102642.

[140] B. Aczel and E. J. Wagenmakers, “Transparency Guidance for ChatGPT Usage in Scientific Writing,” 2023, [Online]. Available: https://psyarxiv.com/b58ex/download?format=pdf.

[141] B. Kutela, K. Msechu, S. Das, and E. Kidando, “Chatgpt’s Scientific Writings: A Case Study on Traffic Safety,” SSRN Electron. J., 2023, doi: 10.2139/ssrn.4329120.

[142] A. Flanagin, K. Bibbins-Domingo, M. Berkwits, and S. L. Christiansen, “Nonhuman ‘authors’ and Implications for the Integrity of Scientific Publication and Medical Knowledge,” JAMA, vol. 329, no. 8, pp. 637–639, 2023, doi: 10.1001/jama.2023.1344.

[143] E. O. Jocelyn Gravel, Madeleine D’Amours-Gravel, “Learning to fake it: limited responses and fabricated references provided by ChatGPT for medical questions,” medRxiv, vol. Pre-print, 2023.

[144] S. P. Thomas, “Grappling with the Implications of ChatGPT for Researchers, Clinicians, and Educators,” Issues in mental health nursing, vol. 44, no. 3. Taylor &Francis, pp. 141–142, 2023, doi: 10.1080/01612840.2023.2180982.

[145] M. Haman and M. Školník, “Using ChatGPT to conduct a literature review,” Account. Res., 2023, doi: 10.1080/08989621.2023.2185514.

[146] Z. Haq, H. Naeem, A. Naeem, F. Iqbal, and D. Zaeem, “Comparing human and artificial intelligence in writing for health journals: an exploratory study,” medRxiv, 2023, doi: 10.1101/2023.02.22.23286322.abstract.

[147] M. P. Polak and D. Morgan, “Extracting Accurate Materials Data from Research Papers with Conversational Language Models and Prompt Engineering –– Example of ChatGPT,” arXiv Prepr*. arXiv2303.05352*, 2023, [Online]. Available: http://arxiv.org/abs/2303.05352.

[148] C. Peter, S. Park, Philipp Schoenegger, “‘Correct answers’ from the psychology of artificial intelligence,” Arxive, 2023.

[149] S. S. Leopold, F. S. Haddad, L. J. Sandell, and …, “Artificial intelligence applications and scholarly publication in orthopaedic surgery,” The Bone &Joint …. boneandjoint.org.uk, 2023, doi: 10.1302/0301-620X.105B.BJJ-2023-0272.

[150] A. Goto and K. Katanoda, “Should we acknowledge ChatGPT as an author?,” Journal of Epidemiology. jstage.jst.go.jp, 2023, [Online]. Available: https://www.jstage.jst.go.jp/article/jea/advpub/0/advpub_JE20230078/_article/-char/ja/.

[151] S. A. Athaluri, S. V. Manthena, V. S. R. K. M. Kesapragada, V. Yarlagadda, T. Dave, and R. T. S. Duddumpudi, “Exploring the Boundaries of Reality: Investigating the Phenomenon of Artificial Intelligence Hallucination in Scientific Writing Through ChatGPT References.,” Cureus, vol. 15, no. 4. cureus.com, p. e37432, 2023, doi: 10.7759/cureus.37432.

[152] H. H. Thorp, “ChatGPT is fun, but not an author,” Science *(80-.).*, vol. 379, no. 6630, p. 313, 2023, doi: 10.1126/science.adg7879.

[153] T. Tate, S. Doroudi, D. Ritchie, and Y. Xu, Educational Research and AI-Generated Writing: Confronting the Coming Tsunami. edarxiv.org, 2023.

[154] N. L. Tenhundfeld, “Two Birds With One Stone: Writing a Paper Entitled ‘ChatGPT as a Tool for Studying Human-AI Interaction in the Wild’ with ChatGPT,” researchgate.net. [Online]. Available: https://www.researchgate.net/profile/Nathan-Tenhundfeld/publication/368300523_Two_Birds_With_One_Stone_Writing_a_Paper_Entitled_ChatGPT_as_a_Tool_for_Studying_Human-AI_Interaction_in_the_Wild_with_ChatGPT/links/63e118c9c97bd76a82765195/Two-Birds-With-One-.

[155] L. Sanchez-Ramos, L. Lin, and …, “Beware of References when Using ChatGPT as a Source of Information to Write Scientific Articles,” Am. J. …, 2023, [Online]. Available: https://pubmed.ncbi.nlm.nih.gov/37031761/.

[156] T. Day, “A Preliminary Investigation of Fake Peer-Reviewed Citations and References Generated by ChatGPT,” Prof. Geogr., 2023, doi: 10.1080/00330124.2023.2190373.

[157] M. Alser and E. Waisberg, “Concerns with the usage of ChatGPT in Academia and Medicine: A viewpoint,” American Journal of Medicine Open. researchgate.net, p. 100036, 2023, doi: 10.1016/j.ajmo.2023.100036.

[158] A. O. Thunström, G. P. Transformer, and S. Steingrimsson, “Does GPT-3 qualify as a co-author of a scientific paper publishable in peer-review journals according to the ICMJE criteria?-A Case Study.” researchsquare.com, 2022, [Online]. Available: https://www.researchsquare.com/article/rs-2404314/latest.pdf.

[159] K. Ide, P. Hawke, and T. Nakayama, “Can ChatGPT be considered an author of a medical article?,” Journal of Epidemiology. jstage.jst.go.jp, 2023, [Online]. Available: https://www.jstage.jst.go.jp/article/jea/advpub/0/advpub_JE20230030/_article/-char/ja/.

[160] J. Y. Lee, “Can an artificial intelligence chatbot be the author of a scholarly article?,” J. Educ. Eval. Health Prof., vol. 20, p. 6, 2023, doi: 10.3352/jeehp.2022.20.6.

[161] N. S. L. Yeo-Teh and B. L. Tang, “NLP systems such as ChatGPT cannot be listed as an author because these cannot fulfill widely adopted authorship criteria.,” Account. Res., Mar. 2023, doi: 10.1080/08989621.2023.2185776.

[162] M. A. Pourhoseingholi, M. R. Hatamnejad, and …, “Does chatGPT (or any other artificial intelligence language tools) deserve to be included in authorship list?,” … Hepatol. from …, 2023, [Online]. Available: https://ojs3.sbmu.ac.ir/ghfbb/index.php/ghfbb/article/view/2747.

[163] M. Polonsky and J. Rotman, “Should Artificial Intelligent (AI) Agents be Your Co-author? Arguments in favour, informed by ChatGPT,” SSRN Electron. J., 2023, doi: 10.2139/ssrn.4349524.

[164] J. A. da Silva, “Is ChatGPT a valid author?,” Nurse Educ. Pract., vol. 68, 2023, doi: 10.1016/j.nepr.2023.103600.

[165] A. Gandhi, P. Satapathy, A. Neyazi, and B. Kumar, “ChatGPT: Roles and boundaries of the new artificial intelligence tool in medical education and health research–Correspondence.” researchgate.net, 2023, [Online]. Available: https://www.researchgate.net/profile/Ahmad-Neyazi/publication/369369666_ChatGPT_Roles_and_boundaries_of_the_new_artificial_intelligence_tool_in_medical_education_and_health_research/links/6424822ea1b72772e4361dac/ChatGPT-Roles-and-boundaries-of-the-new-ar.

[166] S. Yusuf, “ChatGPT, the Blade in Scientific Writing,” Indones. Contemp. Nurs. J., 2023, [Online]. Available: http://journal.unhas.ac.id/index.php/icon/article/view/25634.

[167] R. Rousseau, L. Yang, J. Bollen, and Z. Shen, “Large language models and scientific publishing,” J. Data Inf. Sci., vol. 8, no. 1, p. 1, 2023, doi: 10.2478/jdis-2023-0007.

[168] P. S. Rose and J. S. Fischgrund, “Artificial Intelligence and JAAOS,” … –Journal Am. Acad. …, 2022, [Online]. Available: https://journals.lww.com/jaaos/Fulltext/9900/Artificial_Intelligence_and_JAAOS.669.aspx.

[169] T. Nakaura and S. Naganawa, “Writing medical papers using large-scale language models: a perspective from the Japanese Journal of Radiology,” Jpn. J. Radiol., 2023, doi: 10.1007/s11604-023-01408-z.

[170] J. D. Martínez-Ezquerro, “Authors in the age of language-generation AI: to be or not to be, that is… the question?,” 2023, [Online]. Available: https://osf.io/7uy63/download.

[171] M. R. King, “A Place for Large Language Models in Scientific Publishing, Apart from Credited Authorship,” Cellular and Molecular Bioengineering. Springer, 2023, doi: 10.1007/s12195-023-00765-z.

[172] M. Ali, N. Ahmad, and H. A. Younus, “Integrating chatbots (ChatGPT) in the process of manuscript writing and proposing a roadmap for their future adoption.” authorea.com, 2023, doi: 10.22541/au.168052684.48458398.

[173] N. Anderson et al., “AI did not write this manuscript, or did it? Can we trick the AI text detector into generated texts? The potential future of ChatGPT and AI in Sports & Exercise Medicine manuscript generation.,” BMJ open sport & exercise medicine, vol. 9, no. 1. England, p. e001568, 2023, doi: 10.1136/bmjsem-2023-001568.

[174] Z. N. Khlaif, “Ethical Concerns about Using AI-Generated Text in Scientific Research,” Available SSRN 4387984, 2023, [Online]. Available: https://papers.ssrn.com/sol3/papers.cfm?abstract_id=4387984.

[175] C. M. Hayes, “Did a Robot Write This Title? Creativity, Ownership, Justice, and Copyright Law,” *Creativity, Ownership*, Justice, and Copyright Law …. 2022.

[176] A. Blanco-Gonzalez, A. Cabezon, and …, “The Role of AI in Drug Discovery: Challenges, Opportunities, and Strategies,” arXiv Prepr. arXiv …, 2022, [Online]. Available: https://arxiv.org/abs/2212.08104.

[177] E. Petiska, “ChatGPT cites the most-cited articles and journals, relying solely on Google Scholar’s citation counts. As a result, AI may amplify the Matthew Effect in environmental …,” arXiv Prepr. arXiv2304.06794, 2023, [Online]. Available: https://arxiv.org/abs/2304.06794.

[178] D. Gupta, “The future of Medical Writing: Artificial Intelligence as almost first author and not the ghost,” EFI Bull., 2023, [Online]. Available: https://efi.org.in/journal/index.php/EFIbulletin/article/view/65.

[179] V. Karolinoerita, D. Cahyana, A. Hadiarto, and …, “Application of Chatgpt in Soil Science Research and the Perceptions of Soil Scientists in Indonesia,” *…* Soil Sci. Res. …, [Online]. Available: https://papers.ssrn.com/sol3/papers.cfm?abstract_id=4401008.

[180] J. Wittmann, “Science fact vs science fiction: A ChatGPT immunological review experiment gone awry,” Immunol. Lett., vol. 256–257, pp. 42–47, 2023, doi: 10.1016/j.imlet.2023.04.002.

[181] X. Zhai, “ChatGPT for Next Generation Science Learning,” SSRN Electron. J., 2023, doi: 10.2139/ssrn.4331313.

[182] H. Else, “Abstracts written by ChatGPT fool scientists,” Nature, vol. 613, no. 7944, p. 423, 2023, doi: 10.1038/d41586-023-00056-7.

[183] S. O’Connor and ChatGPT, “Open artificial intelligence platforms in nursing education: Tools for academic progress or abuse?,” Nurse Educ. Pract., vol. 66, p. 103537, Jan. 2023, doi: 10.1016/j.nepr.2022.103537.

[184] Generative Pretrained Transformer, A. O. Thunström, and S. Steingrimsson, “Can GPT-3 write an academic paper on itself, with minimal human input?,” HAL Sci. Ouvert., p. hal-03701250, 2022.

[185] A. Zhavoronkov, “Rapamycin in the context of Pascal’s Wager: generative pre-trained transformer perspective.,” Oncoscience, vol. 9, pp. 82–84, 2022, doi: 10.18632/oncoscience.571.

[186] T. H. Kung et al., “Performance of ChatGPT on USMLE: Potential for AI-assisted medical education using large language models,” PLOS Digit. Heal., vol. 2, no. 2, p. e0000198, 2023, doi: 10.1371/journal.pdig.0000198.

[187] S. O’Connor and ChatGPT, “Open artificial intelligence platforms in nursing education: Tools for academic progress or abuse?,” Nurse Educ. Pract., vol. 66, p. 103537, 2023, doi: https://doi.org/10.1016/j.nepr.2022.103537.

[188] S. O’Connor, “Corrigendum to ‘Open artificial intelligence platforms in nursing education: Tools for academic progress or abuse?’ [Nurse Educ. Pract. 66 (2023) 103537] (Nurse Education in Practice (2023) 66, (S1471595322002517), (10.1016/j.nepr.2022.103537)),” Nurse Educ. Pract., vol. 67, 2023, doi: 10.1016/j.nepr.2023.103572.

[189] B. Kutela, K. Msechu, N. Novat, E. Kidando, and A. Kitali, “Uncovering the Influence of ChatGPT’s Prompts on Scientific Writings using Machine Learning-Based Text Mining Approaches,” SSRN Electron. J., 2023, doi: 10.2139/ssrn.4385895.

[190] “Tools such as ChatGPT threaten transparent science; here are our ground rules for their use,” Nature, vol. 613, no. 7945, p. 612, 2023, doi: 10.1038/d41586-023-00191-1.

[191] J. D. Martínez-Ezquerro, “Authors in the Age of Language-generation AI: To be or not to be, is that Really the Question?,” Arch. Med. Res., 2023, doi: 10.1016/j.arcmed.2023.03.002.

[192] C. Zielinski et al., “Chatbots, ChatGPT, and Scholarly Manuscripts: WAME Recommendations on ChatGPT and Chatbots in Relation to Scholarly Publications,” Open Access Macedonian Journal of Medical Sciences, vol. 11, no. A. aeji.journals.ekb.eg, pp. 83–86, 2023, doi: 10.3889/oamjms.2023.11502.

[193] M. Hammad, “The Impact of Artificial Intelligence (AI) Programs on Writing Scientific Research,” Ann. Biomed. Eng., 2023, doi: 10.1007/s10439-023-03140-1.

